# Comprehensive screening shows that mutations in the known syndromic genes are rare in individuals presenting with hyperinsulinaemic hypoglycaemia

**DOI:** 10.1101/346189

**Authors:** Thomas W Laver, Matthew N Wakeling, Janet Hong Yeow Hua, Jayne AL Houghton, Khalid Hussain, Sian Ellard, Sarah E Flanagan

**Affiliations:** Institute of Biomedical and Clinical Science, University of Exeter Medical School, Exeter, UK; Paediatric Department, Hospital Putrajaya, 62250, Malaysia; Department of Molecular Genetics, Royal Devon and Exeter NHS Foundation Trust, Exeter, UK; Department of Pediatric Medicine, Division of Endocrinology, Sidra Medicine, Doha, Qatar

## Abstract

**Objective:** Hyperinsulinaemic hypoglycaemia (HH) can occur in isolation or more rarely feature as part of a syndrome. Screening for mutations in the ‘syndromic HH’ genes is guided by phenotype with genetic testing used to confirm the clinical diagnosis. As HH can be the presenting feature of a syndrome it is possible that mutations will be missed as these genes are not routinely screened in all newly diagnosed individuals. We investigated the frequency of pathogenic variants in syndromic genes in individuals with HH who had not been clinically diagnosed with a syndromic disorder at referral for genetic testing.

**Design:** We used genome sequencing data to assess the prevalence of mutations in syndromic HH genes in an international cohort of patients with HH of unknown genetic cause.

**Methods:** We undertook genome sequencing in 82 individuals with HH without a clinical diagnosis of a known syndrome at referral for genetic testing. Within this cohort we searched for the genetic aetiologies causing 20 different syndromes where HH had been reported as a feature.

**Results:** We identified a pathogenic *KMT2D* variant in a patient with HH diagnosed at birth, confirming a genetic diagnosis of Kabuki syndrome. Clinical data received following the identification of the mutation highlighted additional features consistent with the genetic diagnosis. Pathogenic variants were not identified in the remainder of the cohort.

**Conclusions:** Pathogenic variants in the syndromic HH genes are rare but should be considered in newly diagnosed individuals as HH may be the presenting feature.

## Introduction

Congenital hyperinsulinaemic hypoglycaemia (HH) is a disorder where episodes of hypoglycaemia are caused by unregulated insulin secretion despite low blood glucose levels. Prompt treatment is crucial in order to avoid serious life-long complications such as seizures and permanent brain injury [1]. Genetic testing is important for the clinical management of this condition as identifying the underlying genetic aetiology will inform on the pancreatic histology which, for patients who are unresponsive to medical treatment, will help to determine whether a lesionectomy or a near-total pancreatectomy is required [2].

HH is most commonly the result of a monogenic aetiology with mutations in at least 8 genes reported to cause isolated disease [1, 3]. Routine screening of these genes using a combination of rapid Sanger sequencing and targeted next generation sequencing identifies a mutation in approximately 40-50% of cases with persistent HH [4]. This suggests that further genetic aetiologies remain to be discovered.

HH has also been reported as a feature in at least 20 different rare genetic syndromes [5]. The most common is Beckwith-Wiedemann syndrome where HH occurs in approximately 50% of cases [6]. More rarely HH has been described in individuals with other overgrowth disorders including Sotos syndrome [7] and growth delay syndromes such as Kabuki [8]. HH has also been reported in some cases with congenital disorders of glycosylation and chromosome abnormalities such as Turner syndrome and Patau syndrome [9-12]. For patients with syndromic HH an early genetic diagnosis is important as this will guide medical management and provide information on prognosis and recurrence risk.

Screening of the syndromic genes is not routinely performed as part of the genetic testing strategy for individuals with HH. The targeted analysis of a gene is usually only performed when there is a clinical suspicion of a specific syndrome in an individual and therefore acts to confirm the clinical diagnosis [13]. Consequently the prevalence of mutations in these genes in HH is not known. As many patients are referred for genetic testing at diagnosis of HH it is possible that some individuals with a mutation in a syndromic gene will not have developed additional extra-pancreatic features at the time of referral for genetic testing. These patients may therefore not receive the most appropriate genetic testing.

In order to assess the frequency of mutations in the known syndromic genes in individuals with HH of unknown genetic cause we performed a comprehensive analysis of genome sequencing data from 82 affected individuals. Screening patients for mutations in the ‘syndromic’ HH genes irrespective of clinical features has the advantage that a genetic diagnosis can precede development of clinical features and guide clinical management, rather than being confirmatory.

## Subjects and methods

### Patient details

82 individuals with HH referred for genetic testing to the Molecular Genetics Laboratory at the Royal Devon and Exeter NHS Foundation Trust were included in the study. All patients had received a clinical diagnosis of HH within the first 12 months of life which was defined by inappropriate insulin or C-peptide secretion and/or inappropriately low plasma β-hydroxybutarate and free fatty acids at the time of hypoglycaemia. In all cases the HH had persisted for greater than 6 months or had required pancreatic resection following a poor response to treatment. In 15 patients extra-pancreatic features were present at the time of referral for genetic testing. None of these patients had received a clinical diagnosis consistent with a known syndromic form of HH at the time of study. Mutations in the *ABCC8*, *KCNJ11*, *HADH*, *HNF4A*, *HNF1A*, *GLUD1*, *GCK* and *SLC16A1* genes had been excluded in all patients using targeted next generation sequencing [13].

### Gene panel and variant calling

We utilised the Phenomizer bowser to identify disease entries annotated for hyperinsulinaemic hypoglycaemia (HPO id: 0000825) [14]. A survey of the literature was also undertaken to identify further genes in which mutations have been reported to cause HH as part of a syndrome.

Whole genome sequencing was performed on DNA extracted from peripheral blood leukocytes of the 82 probands. All samples were sequenced on an Illumina HiSeq 2500 or Illumina X10 with a mean read depth of 33.25 (standard deviation 4.25). The sequencing data was analysed using an approach based on the GATK best practice guidelines [15]. This involved aligning the reads to the hg19/GRCh37 human reference genome with BWA mem [16], applying Picard for duplicates removal [17], and GATK IndelRealigner for local re-alignment and running the base quality score realignment [18]. GATK haplotypecaller was used to identify variants which were annotated using Alamut batch version 1.8 (Interactive Biosoftware, Rouen, France) and variants which failed the QD2 VCF filter or had less than 5 reads supporting the variant allele were excluded. Variants that passed filtering were confirmed by Sanger sequencing and tested in the parents (details of primer sequences are available on request). CNVs were called by SavvyCNV [19] and runs of common single nucleotide polymorphisms were analysed to investigate uniparental isodisomy.

## Results

We identified 20 genetic syndromes in which HH has been reported as a feature (Table 1). In 82 patients we screened the coding regions and intron/exon boundaries of the 18 genes associated with these syndromes. We also searched for copy number variations (CNVs) and evidence of uniparental isodisomy at genomic regions associated with Beckwith-Wiedemann, Patau and Turner syndromes.

**Table 1:**
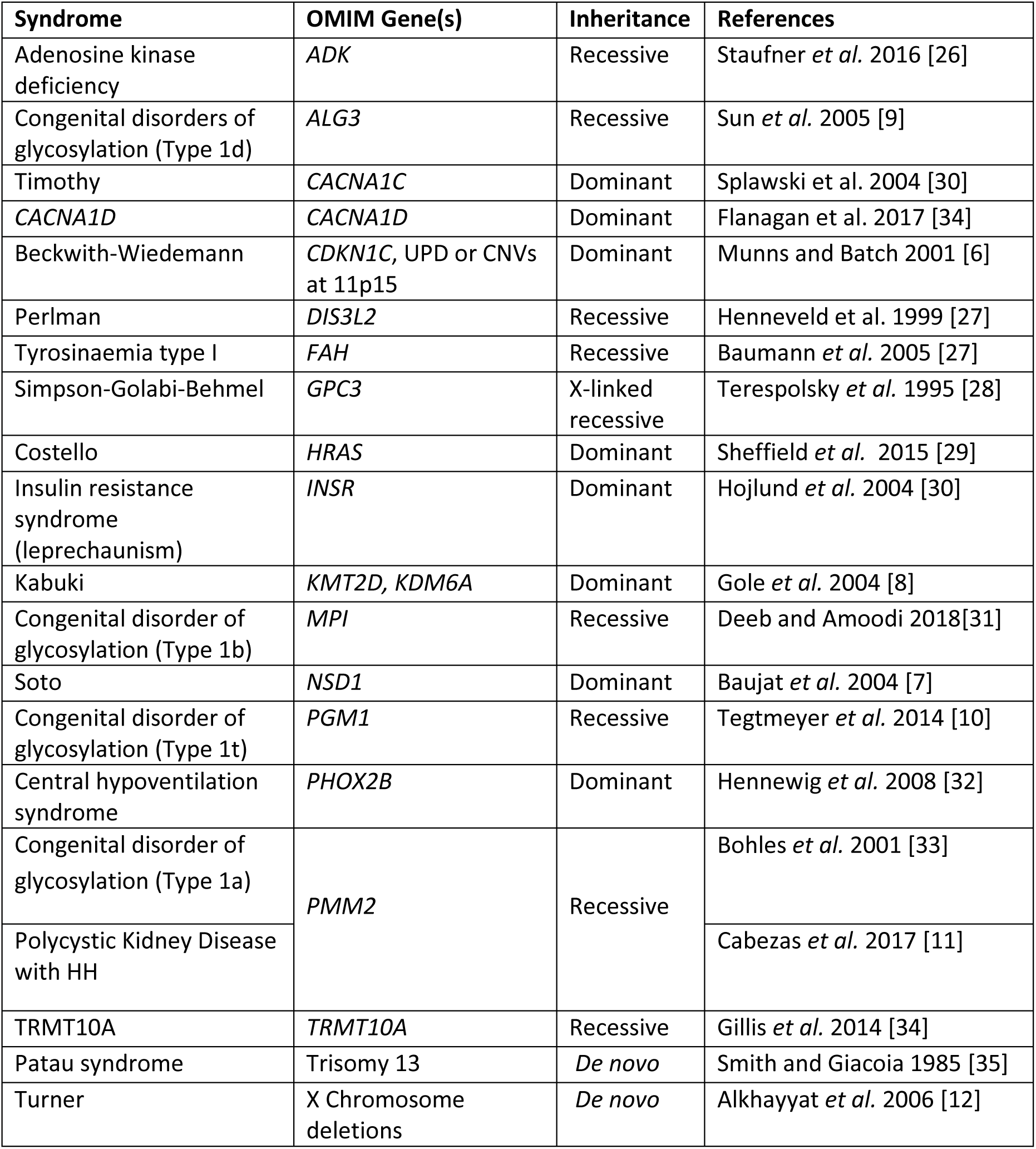
Syndromes in which Hyperinsulinaemic Hypoglycaemia (HH) has been reported as a feature. The 18 genes in which mutations have been reported to cause syndromic HH plus the three genomic regions known to be affected by copy number variants (CNVs) or uniparental isodisomy (UPD) are provided.

We identified 70 non-synonymous (nonsense, frameshift, splice site, missense) variants in the 18 genes. As the incidence of HH in outbred populations is estimated to be between 1 in 27,000 [20] and 1 in 50,000 [21] we excluded all variants that were present in gnomAD controls [22] at a frequency greater than 1 in 27,000. In addition, we excluded variants which did not fit the known inheritance pattern of the syndrome: excluding single variants in recessive genes and variants inherited from an unaffected parent in dominant genes. This left one *de novo* variant in the *KMT2D* gene where dominantly inherited loss of function mutations are reported to cause Kabuki syndrome[8].

CNVs were not detected at any of the regions analysed and analysis of single polymorphisms excluded uniparental isodisomy at the Beckwith Wiedemann syndrome locus at chromosome 11p15.5. All patients had normal dosage of chromosome 13 and all females had a 46XX karyotype which excluded Patau and Turner syndrome respectively.

The heterozygous frameshift variant in the *KMT2D* gene p.(Lys5244Serfs*13), (c.15731_15732del) (NM_003482.3) had arisen *de novo* in the proband and was classified as ‘Pathogenic’ according to the American College of Medical Genetic guidelines [23]. The female patient was of Sarawakian ethnicity born in Malaysia to non-consanguineous parents at 38 weeks gestation with a birth weight of 3.36kg. During the pregnancy her mother was diagnosed with gestational diabetes that was controlled by diet and the pregnancy was complicated by polyhydraminos. Hypoglycaemia was diagnosed in the proband at birth (2.0mmol/l) which required intermittent intravenous dextrose for recurrent symptomatic hypoglycaemia until day 30 of life. Diazoxide treatment (5 mg/kg/day) and hydrochlorothiazide (1.5 mg/kg/ twice daily) was started at 30 days which resulted in euglycaemia. Hyperinsulinism as the cause of hypoglycaemia was confirmed biochemically. The patient required continuous diazoxide treatment and a nasogastric tube for feeding. At the age of 6 months the patient was referred for genetic testing for persistent HH. At this point the only extra-pancreatic feature reported to the genetics laboratory was gastro-oesophageal reflux. At follow up subsequent to the genetic diagnosis clinical features consistent with a diagnosis of Kabuki syndrome were reported. At the age of 5 months her weight was 6.06 kg (10^th^-50^th^ centile), height was 60 cm (<10^th^ centile) and head circumference was 38.5 cm (<10^th^ centile). She had soft facial dysmorphism, hypotonia and laryngomalacia with inspiratory stridor. Her gross and fine motor development was delayed. She also had a resolved small muscular ventricular septal defect and congenital small right kidney with normal left kidney. She had recurrent pneumonia and died before reaching 1 year of age.

## Discussion

We screened for genetic aetiologies where HH has been reported to feature as part of a complex syndrome in 82 patients with HH of unknown cause. We identified one patient with a pathogenic variant in *KMT2D* that confirmed a diagnosis of Kabuki syndrome [24]. At referral the patient was reported to the genetics laboratory as having persistent HH and gastro oesophageal reflux. At follow up subsequent to the genetic diagnosis clinical features consistent with a diagnosis of Kabuki syndrome were reported. These included a structural heart defect with growth and developmental delay. As the patient died before the age of 1 year it is not possible to ascertain if they would have developed further features of this syndrome.

Mutations in the ‘syndromic’ HH genes were not identified in the majority of our cohort (81/82 patients). This low pick-up rate is likely to reflect referral bias. Our patients were selected due to the presence of persistent HH rather than a syndromic phenotype. It is likely that for the majority of cases with syndromic disease additional features are evident from birth allowing for a clinical diagnosis followed by confirmatory genetic testing. This would consequently reduce the prevalence of cases within our cohort as the majority of patients are referred for routine screening of the 8 known isolated HH genes. We can also not rule out the possibility that some patients have mutations in other genes known to cause multi-system disease which were not screened in this study as HH has not been recognised as a common feature. Furthermore it is possible that some patients with Beckwith-Wiedemann syndrome have a methylation defect in the absence of a structural abnormality that was not detected by our analysis.

The finding that the patient with Kabuki syndrome had HH as the presenting feature highlights the need to consider screening for the syndromic genes in individuals newly diagnosed with HH. Identifying the underlying genetic aetiology is important for these patients as it will inform on prognosis which will allow for better clinical management. A genetic diagnosis also provides important information on recurrence risk.

Although we found that mutations causing these 20 syndromes are rare, the recent adoption of targeted next-generation sequencing by molecular genetic laboratories makes screening of multiple genes for conditions such as HH feasible. Whilst there are some disadvantages of this approach in terms of the difficulties with variant interpretation in the absence of additional clinical features, the benefits that an early and an accurate genetic diagnosis has on patient care outweigh the extra-time required for the analysis of the genetic data. It is also anticipated that variant interpretation will become easier through data sharing initiatives such as ClinVar [25] and as the number of publically available control datasets increase, enabling more variants to be excluded by frequency [22].

Screening large cohorts for mutations in these genes will provide further information on the prevalence of mutations in HH which will help to guide future genetic screening strategies for this condition and may also identify individuals with ‘non-classical’ features which would expand the phenotype associated with these syndromes. Furthermore, systematic screening of these genes would also prevent individuals with a known cause of HH from being included in costly gene discovery studies.

In conclusion, although mutations in the syndromic HH genes are rare in individuals referred for routine testing for HH, the clinical impact of finding a mutation and availability of targeted next generation sequencing means that inclusion of these genes on panels should be considered by molecular genetic laboratories routinely screening for this condition.

## Author contributions

TWL and SEF designed the study. JHYH and KH recruited patients and performed clinical phenotyping. TWL and MNW analysed the sequencing data. JALH, SE and SEF performed variant interpretation. TWL and SEF wrote the manuscript. All authors reviewed and approved the manuscript.

## Acknowledgements

The authors thank Rebecca Ward (University of Exeter, Exeter, UK) for her technical assistance.

## Figure legends

**Figure 1:**
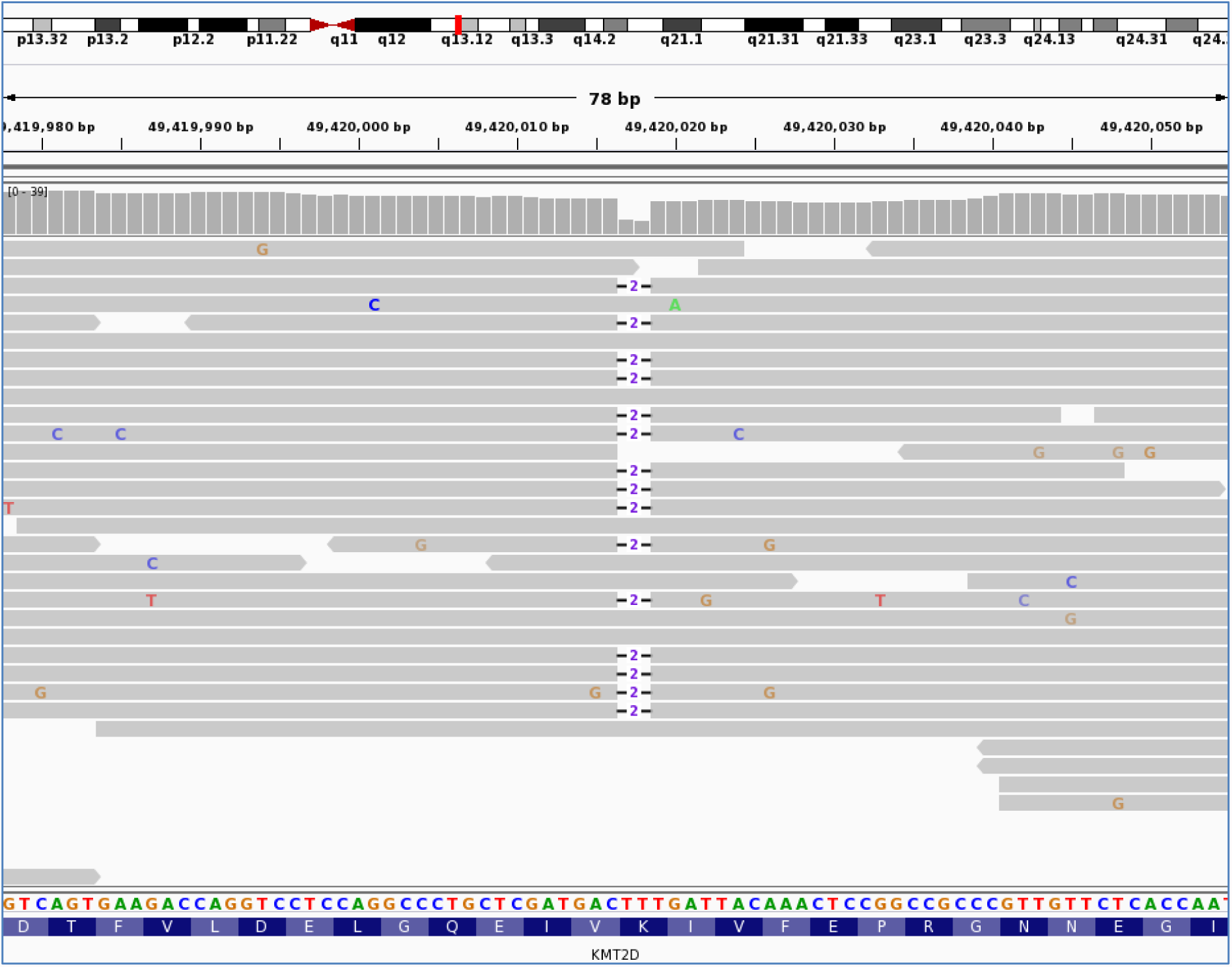
*KMT2D* variant g.49420017_49420018del/c.15731_15732del/p.Lys5244Serfs*13. Visualised in integrative genomics viewer (IGV). It shows the sequencing reads (grey bars) mapping to exon 48 of the *KMT2D* gene located at genomic position g. 49,420,017 on chromosome 12. The reference nucleotide sequence and the amino acid translation are provided under the sequencing reads. The heterozygous deletion of TT is illustrated by -2- and is present in 15 of the 25 sequencing reads present at this position. The deletion causes a frameshift.

